# Functional diversity metrics using kernel density *n*-dimensional hypervolumes

**DOI:** 10.1101/2020.01.25.919373

**Authors:** Stefano Mammola, Pedro Cardoso

## Abstract

1. The use of *n*-dimensional hypervolumes in trait-based ecology is rapidly increasing. By representing the functional space of a species or community as a Hutchinsonian niche, the abstract Euclidean space defined by a set of independent axes corresponding to individuals or species traits, these multidimensional techniques show great potential for the advance of functional ecology theory.
2. In the panorama of existing methods for delineating multidimensional spaces, the R package *hypervolume* [*Glob. Ecol. Biogeogr.* (2014) 23:595–609] is currently the most used. However, functions for calculating the standard set of functional diversity (FD) indices—richness, divergence, and regularity—have not been developed within the *hypervolume* framework yet. This gap is delaying its full exploitation in functional ecology, meanwhile preventing the possibility to compare its performance with that of other methods.
3. We develop a set of functions to calculate FD indices based on *n*-dimensional hypervolumes, including alpha (richness), beta (and respective components), dispersion, evenness, contribution, and originality. Altogether, these indices provide a coherent framework to explore the primary mathematical components of FD within a multidimensional setting. These new functions can work either with hypervolume objects or raw data (species presence or abundance and their traits) as input data, and are versatile in terms of input parameters and options.
4. These functions are implemented within *BAT* (Biodiversity Assessment Tools), an R package for biodiversity assessments. As a coherent corpus of functional indices based on a common algorithm, it opens the possibility to fully explore the strengths of the Hutchinsonian niche concept in community ecology research.

## Introduction

The past two decades have seen an increasing interest on different facets of biodiversity, namely on the way species interact among them and with their environment, with the resulting exponential growth of functional ecology studies. This push was primarily driven by a few milestone theoretical contributions illustrating a way of reshaping entire areas in ecology from a functional perspective, including population and community ecology (Petchey & Gaston, 2002; McGill, Enquist, Weiher, & Westoby, 2006; Bolnick et al., 2011; Violle et al., 2012), conservation biology (Wellnitz & Poff, 2001; Cadotte, Carscadden, & Mirotchnick, 2011), and biogeography (Violle, Reich, Pacala, Enquist, & Kattge, 2014). This fast development of functional ecology theory has led to a proliferation of mathematical approaches for estimating and visualizing functional diversity (FD) (Legras, Loiseau, & Gaertner, 2018)—raw data, distance matrices, trees, hypervolumes, etc. (Figure 1)—as well as indices for measuring different mathematical facets underlying FD (Mouchet, Villéger, Mason, & Mouillot, 2010; Schleuter, Daufresne, Massol, & Argillier, 2010; Guillerme, Puttick, Marcy, & Weisbecker, 2019).

**Figure 1.**
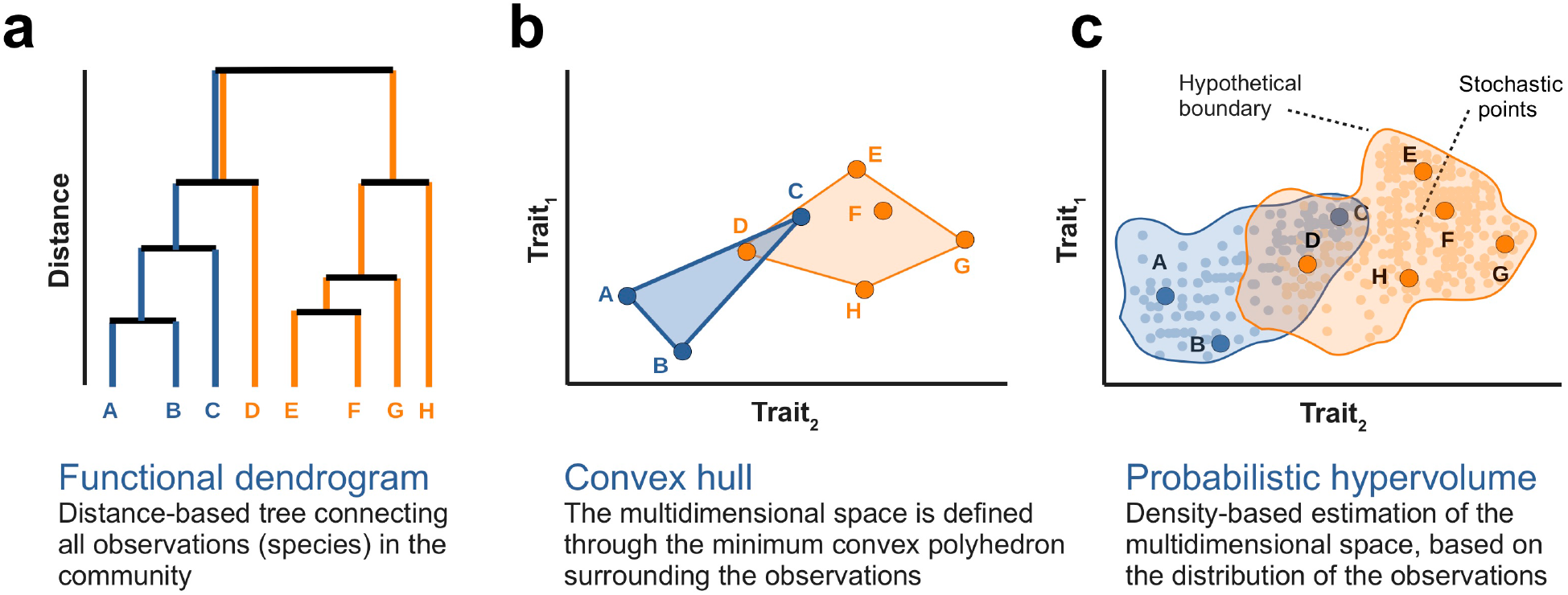
Three examples of mathematical representations of functional diversity (FD) of communities. Examples are based on two hypothetical traits and two communities, represented in turquoise and orange. a) The trait space can be represented as a functional dendrogram (=tree), whose construction is based on a trait-based distance matrix among species. Using this framework, the functional richness of a community can be estimated as the total branch length of a tree linking all species represented in such community—in the example, the FD of a community with 3 (in turquoise) and 5 species (in orange) are shown. b) The trait space can be represented as the minimum convex hull comprising the species occupying the trait space. The functional richness of a community can be estimated as the area of the convex hull. c) The trait space can be constructed using kernel density hypervolumes, whereby it is approximated as a cloud of stochastic points sampled based on a set of observations (e.g. the traits of the species in the community). The functional richness of the community can be estimated as the volume of the hypervolume delineated by the stochastic points.

In the early 2000s, probably the most popular visualization of FD was the functional tree (Figure 1 a), the distance-based dendrogram connecting all functional elements in the community (Petchey & Gaston, 2002, 2006). Nevertheless, given that trait data are multivariate in essence, there have been a recent upsurge of methods relying on raw position of species or individuals within a multi-dimensional space (Guillerme et al., 2019), without transforming data into dissimilarities. The simpler approach is the convex hull hypervolume (Figure 1b), introduced by Cornwell et al. (2006) and later popularized by the R package *FD* (Laliberté, Legendre, & Shipley, 2014). This is a widely used and computationally fast method; yet, it has some key limitations (Podani, 2009), especially the assumption that there is no empty space within extreme values of traits (Blonder, 2016). In response to these limitations, a new family of probabilistic hypervolumes (Figure 1c) have since been developed, based on kernel density estimations (Blonder, Lamanna, Violle, & Enquist, 2014; Blonder et al., 2018), multivariate normal distributions (Swanson et al., 2015), dynamic range boxes (Junker, Kuppler, Bathke, Schreyer, & Trutschnig, 2016), or trait probability densities (Carmona, de Bello, Mason, & Lepš, 2016a, 2016b, 2019), among others.

The *hypervolume* R package (Blonder, 2018) is the most popular among these novel probabilistic approaches; it uses high-dimensional kernel density estimations to delineate the shape and volume of the multi-dimensional space (Blonder et al., 2014, 2018). Other packages based on multidimensional spaces, such as *FD* (Laliberté et al., 2014) or *TPD* (Carmona et al., 2019), include a vast collection of indices that allow the user to explore a functional community based on different mathematical aspects of FD, namely the richness, divergence, and regularity components (Pavoine & Bonsall, 2011; Tucker et al., 2017). Conversely, *hypervolume* was not explicitly developed for functional analyses—for example, it is often used to explore bioclimatic niches (Blonder, Lamanna, Violle, & Enquist, 2017)—and as yet, analogous FD indices have not been developed. This gap is hindering its use in trait-based ecology and is preventing the possibility to fully compare its performance with that of alternative methods.

Here, we describe a new set of R functions (Table 1) to calculate the analogous of standard functional metrics with kernel density *n*-dimensional hypervolumes. These functions are made available through *BAT* (Biodiversity Assessment Tools) (Cardoso, Mammola, Rigal, & Carvalho, 2020), an R package that provides an assortment of functions for exploring taxonomic (TD), phylogenetic (PD), and functional (FD) components of biological diversity (Cardoso, Rigal, & Carvalho, 2015).

**Table 1.** Classification of the new functional indices using the Pavoine–Bonsall scheme. The row entries distinguish between indices calculated at the observation level or hypervolume levels (within and between), while the column entries represent the three dimensions of Richness, Divergence, and Regularity (Pavoine & Bonsall, 2011; Tucker et al., 2017). n/a = not available

|  |  | **FUNCTIONAL DIVERSITY COMPONENT**<br>[following the Pavoine–Bonsall (2011) scheme for classifying biodiversity metrics] |  |  |
| --- | --- | --- | --- | --- |
|  |  | **RICHNESS**<br>(indices reflecting the sum of difference among observations) | **DIVERGENCE**<br>(indices reflecting the average difference among observations) | **REGULARITY**<br>(indices reflecting how regular the difference among observations are) |
| **Observation level**<br>(species within community, individuals of a species, etc.) | **Definition** | How unique is an observation within the total trait space | Dissimilarity between observations within the total trait space | n/a |
|  | **Calculation** | Net contribution of the observation to the total volume of the hypervolume | Average dissimilarity between the observation and a sample of random points within the boundaries of the hypervolume |  |
|  | **R function** | `kernel.contribution` | `kernel.originality` |  |
| **Hypervolume level within**<br>(properties of the trait space of a species or a community) | **Definition** | Total amount of trait space available | How spread and dense is the trait space | How regular is the trait space |
|  | **Calculation** | Volume of the hypervolume | Average difference between the trait space centroid and random points within the boundaries of the hypervolume | Overlap between the observed hypervolume and a theoretical, perfectly even hypervolume |
|  | **R function** | `kernel.alpha` | `kernel.dispersion` | `kernel.evenness` |
| **Hypervolume level between**<br>(between species or communities) | **Definition** | How dissimilar are two or more trait spaces | n/a | n/a |
|  | **Calculation** | Replacement and net difference in amplitude components of hypervolumes (*kernel.beta*) or other measures of centroid distance, intersection, or overlap (*kernel.similarity*) |  |  |
|  | **R function** | `kernel.beta`<br>`kernel.similarity` |  |  |

### New functionalities in BAT

#### INPUT DATA AND DOMAIN OF APPLICABILITY

The new functions presented here (Figure 2) represent the analogous of the functional indices that were originally developed in *BAT* for phylogenetic and functional trees (Cardoso et al., 2015). The calculation of the new indices based on hypervolumes requires the user to input either:

i. a `‘Hypervolume’` class object;
ii. a `‘HypervolumesList’` class object; or
iii. raw data matrices with species composition of the communities and their functional traits.

Objects of class `‘Hypervolume’` and `‘HypervolumeList’` can be constructed outside the *BAT* environment, using the function `hypervolume` in the *hypervolume* package (Blonder, 2018). Alternatively, hypervolumes can be constructed directly within *BAT* by feeding the functions with raw data, namely a sites x species matrix with incidence or abundance data about the species in the community and a species x traits matrix (or individuals if species-level analyses are done). If abundance data are used (`abund = TRUE`), each observation is weighted by replicating it times the abundance in the estimation of the hypervolume.

**Figure 2.**
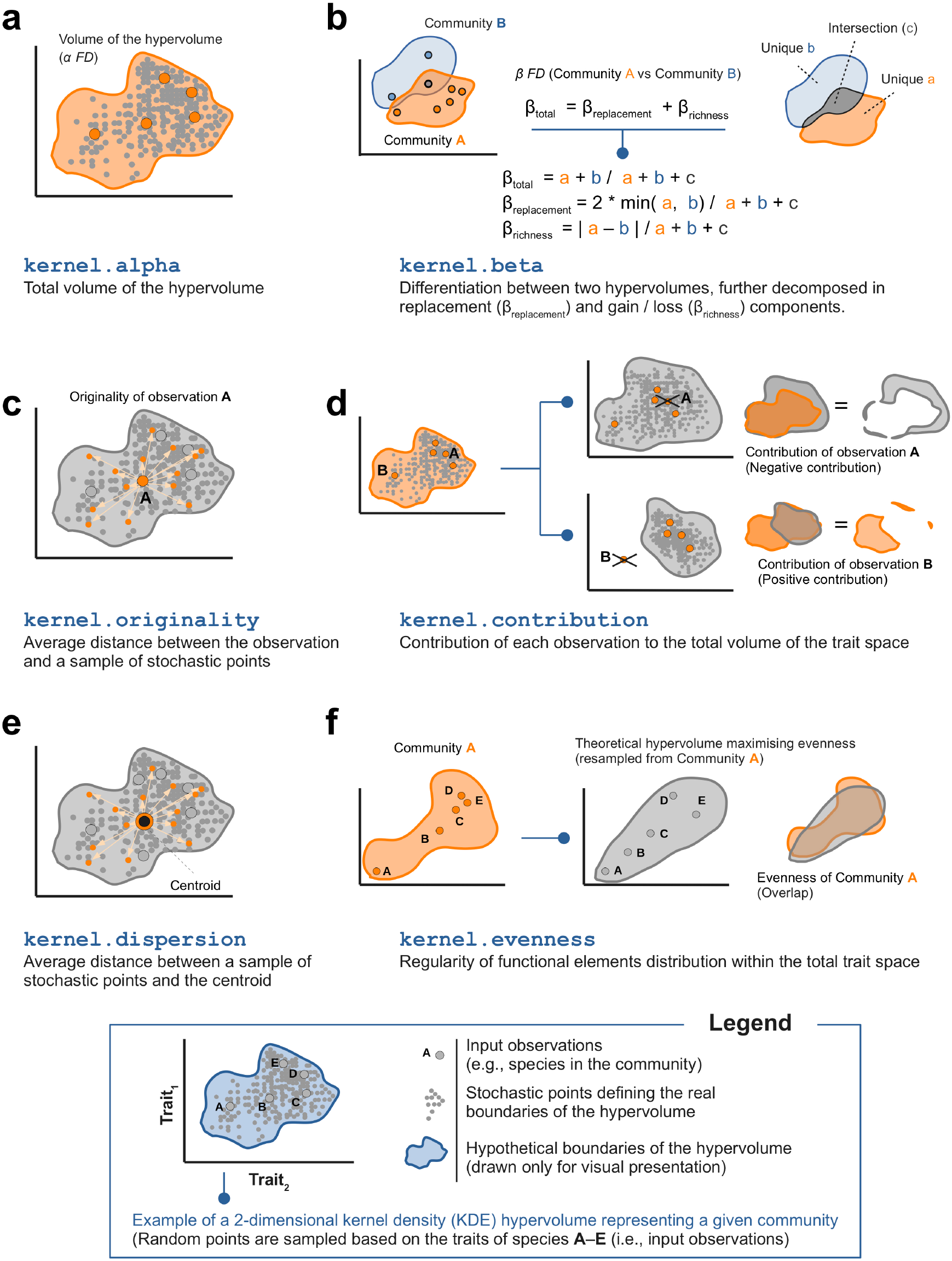
Graphical illustration of each new functions based on hypothetical 2-dimensional hypervolumes representing functional communities. **a**) `kernel.alpha` measures functional richness as the total volume of the hypervolume. **b**) `kernel.beta` measures functional dissimilarity between hypervolumes using the β_total_ = β_replacement_ + β_richness_ partition framework (Carvalho & Cardoso, 2018). **c**) `kernel.originality` is the average distance of the observation and a sample of stochastic points constituting the hypervolume, thus measuring how unique is the position of individuals observations in the trait space. **d**) `kernel.contribution` measures the contribution of each observation to the total volume of the trait space using a leave-one-out approach. In this example, the contribution of A is negative, because the observation is clustered with other observations, which leads a higher proportion of stochastic points to cluster in the vicinity of these observations. By removing A, the stochastic points are “freer” to spread in the trait space, which results in a slight increase in the volume. Instead, the contribution of B is positive, since the hypervolume constructed without this observations is less voluminous than the initial one. **e**) `kernel.dispersion` measures how spread is the trait space, and is estimated either as the average distance between a sample of stochastic points and the hypervolume centroid or, alternatively, as the average distance between a sample of stochastic points. **f**) `kernel.evenness` measures the regularity of stochastic points distribution within the total trait space. Evenness is calculated as the overlap between the input hypervolume and a second, imaginary hypervolume where traits and abundances are evenly distributed within their possible range (Carmona et al., 2016b).

Stochastic points determining the shape and volume of the hypervolume (Figure 1c) are estimated in *hypervolume* using three alternative methods (Blonder et al., 2018): high-dimensional kernel density estimation (KDE) (`method = “gaussian”`) (Blonder et al., 2014), support vector machine (SVM) delineation (`method = “SVM”`), and convex hull generation procedure (`method = “box”`) (Blonder et al., 2018). Currently BAT supports all methods, but we recommend KDE hypervolumes (`“gaussian”`), because other methods assumes the probability density to be constant throughout the distribution. The convex hull delineation (`“box”`) provides a representation of the trait space similar to that of the package *FD*. Since this method is computationally faster than the others, it can be useful for a quick explorative screening of the trait space. The properties of SVM hypervolumes have not been studied yet.

Two or more hypervolumes can only be compared if they are constructed using the same number (and type) of traits. Traits should either not be highly correlated (threshold at Pearson |*r*|≥ 0.8; Blonder, 2018), or transformed through different means (e.g. PCoA) so to have orthogonal hypervolume axes. Although traits should preferentially be continuous variables, transformation also allows implementing categorical traits (Carvalho & Cardoso, 2018).

While the new *BAT* functions have been specifically developed for trait-based hypervolumes, these can also be applied to other types of input data, for example to explore bioclimatic niches or behavioral hypervolumes.

#### BEST PRACTICE & RECOMMENDATION

Given that the density and positions of stochastic points in the trait space are probabilistic, the output of each FD index will intimately depend on the quality of input hypervolumes. Important factors and parameters to consider are:

i. if enough observations are available for hypervolume delineation;
ii. the algorithm to be used for hypervolume construction; and
iii. the choice of bandwidths and other input parameters.

We refer the reader to Blonder et al. (2014, 2018) for details on these features.

The number of traits used for hypervolume delineation is a critical feature to consider as well, directly translating in the number of dimensions (= axes). As dimensionality gets high, a multidimensional space is subject to the “curse of dimensionality” (Bellman, 1957), whereby stochastic points will become sparser and will “migrate” towards the hypervolume boundaries (Blonder, 2016; Guillerme et al., 2019; Mammola, 2019). Also, in high dimensional settings, the probability of overlap between any two hypervolumes will decrease (Mammola, 2019), thus affecting the estimation of dissimilarity indices (i.e., functions `kernel.beta` and `kernel.similarity`). Insofar as computation time also scale exponentially with dimensionality, whenever possible we recommend keeping the number of dimensions low (e.g., 3– 5).

When comparing multiple communities in terms of FD, we recommend to follow best practice for assessing the similarity of *n*-dimensional hypervolumes (Mammola, 2019).

#### FUNCTION DESCRIPTION

##### kernel.alpha

This function calculates functional richness of the trait space (Figure 2a). Richness is simply the total volume of the functional hyperspace, as returned by the function `get_volume` (Blonder, 2018).

##### kernel.beta

This function estimates functional β-diversity based on the framework recently proposed by Carvalho & Cardoso (2018), computing a pairwise decomposition of the overall differentiation among kernel hypervolumes (β_total_) into two components: the replacement of space between hypervolumes (β_replacement_), and gain/loss of space enclosed by each hypervolume (β_richness_). In other words, this approach decomposes β-diversity into its drivers in a similar way as already proposed for functional trees (Cardoso et al., 2014). Each β component can range from 0 (when hypervolumes are identical) to 1 (fully disjunct hypervolumes), and β_total_ = β_replacement_ + β_richness_. See formula in Figure 2b and Cardoso & Carvalho (2018) for more details. This decomposition has several advantages over other options that do not reflect the true drivers of beta diversity (see Carvalho, Cardoso, Borges, Schmera, & Podani, 2013 for a discussion).

##### kernel.originality

A measure of the originality of each observation used to construct the hypervolume (Pavoine, Ollier, & Dufour, 2005), calculated as the average distance between each observation to a sample of stochastic points within the boundaries of the hypervolume (Figure 2c). The number of sample points to be used in the estimation is controlled by the frac parameter. By setting `relative = TRUE`, the originality values are returned relative to the most original species (which will take the value of 1).

##### kernel.contribution

This function evaluates the contribution of each observation to the total volume of the hypervolume (Figure 2d). It does so via a leave-one-out approach, whereby the contribution of each observation is calculated as the difference in volume between the full hypervolume and a second hypervolume constructed without that specific observation. Note that, interestingly, the contribution of an individual observation could be negative, if the removal of this observation increases the volume of the total hypervolume. This might happen, although not always, in cases when the presence of a given species decreases the average distance between all the species in the community, i.e., when a given species is close to the “average” species of that community, making that community less diverse in some sense. This does not happen in the case of functional dendrograms or convex hulls.

##### kernel.dispersion

Hypervolume dispersion (Figure 2e) is calculated as the average distance between a sample of stochastic points and the centroid (Laliberté & Legendre, 2010), thus making a close analogy with the implementation of divergence indices in *FD* (Laliberté et al., 2014) or *TPD* (Carmona et al., 2019). An alternative calculation is also implemented (`func = ‘dissimilarity’`), whereby dispersion is expressed as the average pairwise distance of a sample of stochastic points in the trait space. This makes the closest analogy with the dispersion of a phylogenetic or functional tree (function `dispersion`; Cardoso et al., 2015). In both cases, the number of stochastic points used in the estimation is controlled by the `frac` parameter.

##### kernel.evenness

This function measures the functional evenness of an hypervolume, namely the regularity of functional elements distribution within the total trait space (Mason, Mouillot, Lee, & Wilson, 2005; Villéger, Mason, & Mouillot, 2008). Following a similar approach to that of Carmona et al. (2016b, 2019), evenness is calculated as the overlap between the input hypervolume and a second, imaginary hypervolume where traits and abundances are evenly distributed within their possible range (Figure 2f).

##### kernel.similarity

A versatile function to compare multiple hypervolumes, by calculating their pairwise centroid distance, minimum distance, intersection, and Jaccard and Sørensen-Dice similarity (see Mammola, 2019 for details).

### Example analysis

To illustrate the new functions, we provide an example analysis based on a study of ground-dwelling arthropods in sampling plots characterized by different levels of urbanization within the municipality of Turin (NW-Italy) (Piano, Giuliano, & Isaia, 2020). In order to test the effects of patch isolation within the urban matrix on the FD of arthropod assemblages, the authors identified two types of subplots within each plot, where they used pitfall traps to sample ground-dwelling fauna. In particular:

i. isolated subplots are patches within a traffic roundabout, surrounded by roads; and
ii. connected subplots are patches in green areas, connected with the surrounding environmental matrix.

We extracted from this database a random subset of 60 subplots, 30 isolated and 30 connected. For each subplot, we took abundance data for the spiders’ community and three traits for each species: body length, dispersal strategy, and hunting strategy. Categorization of hunting strategies was based on Cardoso et al. (2011), whereas three type of dispersal (1= non- or sporadic ballooners; 2= ballooners as juveniles; 3= ballooners at all life stages) were assigned by Piano et al. (2020) based on literature (Bell, Bohan, Shaw, & Weyman, 2005; Blandenier, 2009; Simonneau, Courtial, & Pétillon, 2016).

We used the new functions to explore FD of spider communities in isolated versus connected subplots, using *BAT* version 2.0.1. (Cardoso et al., 2020) and *hypervolume* version 2.0.11. (Blonder, 2018). Since spiders’ functional traits included two categorical traits, we followed the approach proposed by Carvalho & Cardoso (2018) to use categorical variables with hypervolumes. We applied a Gower dissimilarity measure (Gower, 1971) to the complete trait matrix and then, we analyzed the resulting distance matrix through Principal Coordinate Analysis (PCoA) in order to extract orthogonal axes for hypervolume delineation. We retained the first three PCoA axes (cumulatively explaining 95.5% of the total variance) to construct hypervolumes, using the gaussian method and a default bandwidth (Blonder et al., 2018). We then estimated functional richness of each community with `kernel.alpha`.

On average, connected subplots had a slightly higher functional richness (Figure 3a). In fact, connected subplots are more easily colonized by a larger number of spider species, that in turn accounts for a greater diversity of traits. Isolated subplots, on the other hand, are mostly colonized by a subset of specialized species possessing specific dispersal traits (smaller body size, greater dispersal ability) allowing them to reach these isolated habitats. Traits of spider communities in connected subplots were not significantly more dispersed (`kernel.dispersion`, calculated with the `func = ‘divergence’`; Figure 3b) or homogeneous (`kernel.evenness`; Figure 3c) than those in isolated subplots (Dispersion: t-test= 1.25, df= 44.94, p= 0.22; Evenness: t-test= – 0.13, df= 57.04, p= 0.89).

**Figure 3.**
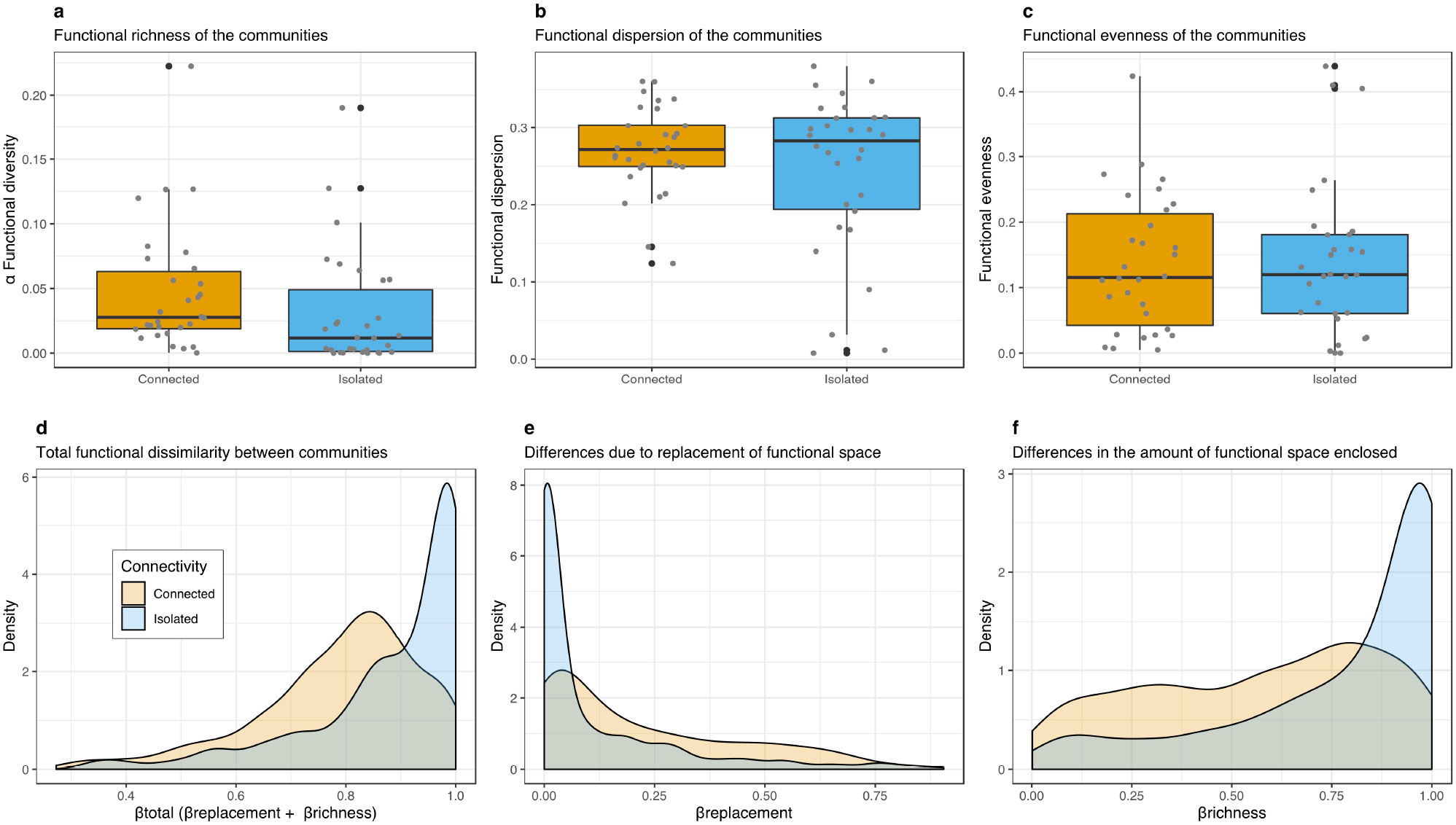
Results of the example analysis based on Piano et al. (2020) dataset. **a–c**) Functional richness, dispersion, and evenness of spider communities in connected and isolated subplots. **d**–**f**) Density of β functional diversity values for pairwise comparison of communities in connected and isolated subplots. Total functional β diversity (β_total_) is split in two components: β_replacement_ is turnover in functional composition explained by replacement of species alone, and β_richness_ is the turnover in functional composition explained by species loss/gain alone.

We explored dissimilarities (*β*) in traits among communities inhabiting the two different types of subplots with the function `kernel.beta`. β_total_ values were higher in pairwise comparisons between communities in isolated subplots, whereas communities in connected subplots were in general more similar to one another (Figure 4d). This result suggest that colonization of traffic roundabout is less predictable and more subject to higher differences in richness:: only a subset of spiders with higher dispersal potential and probably higher resistance to disturbed habitats are able to reach and survive in isolated subplots. This is further confirmed by the lower values of β_replacement_ (Figure 3e) and higher values of β_richness_ (Figure 3f) in isolated subplots.

**Figure 4.**
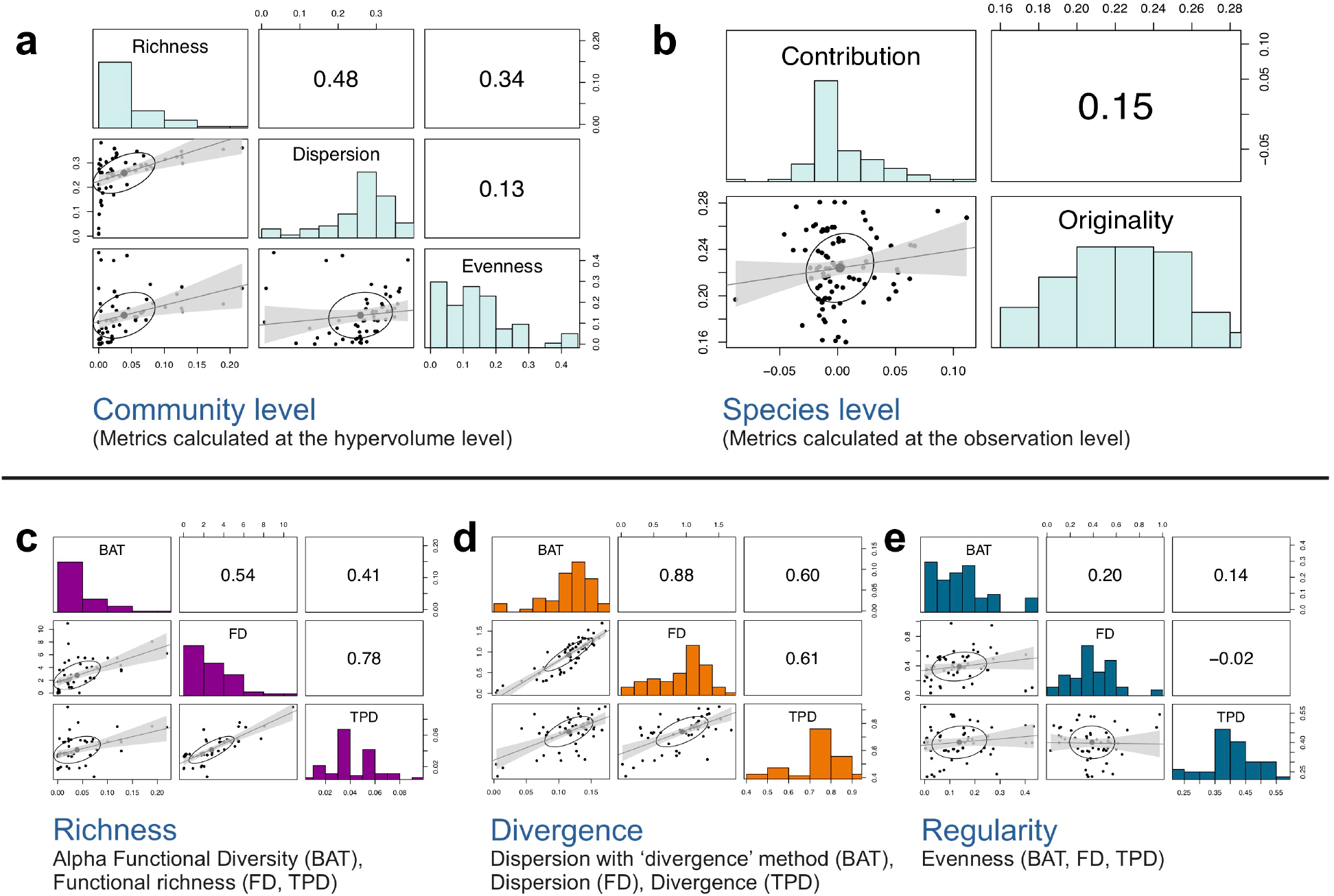
Pairwise Pearson’s correlations among the indices calculated within BAT and with other packages for functional diversity analyses in a multidimensional space (FD, TPD). **a**–**b**) Correlations among indices calculated at the hypervolume (**a**) and observation (**b)** levels. **c**–**e**) Correlations among functional richness (**c**), divergence/dispersion (**d**) and evenness (**e**) calculated with the package BAT, FD, and TPD. The analyses are based on the dataset by Piano et al. (2020). For each panel plot, histograms on the diagonal display the distribution of values. Bivariate scatter plots are displayed below the diagonal and the Pearson *r* correlations above the diagonal. For each scatter plot, regression line and correlation ellipse are included for visual presentation.

By using the function `kernel.contribution`, we further inferred that *Nurscia albomaculata* (Lucas, 1846) (Titanoecidae) was the species contributing the most to the trait space in isolated subplots, being the only species of large size (>10 mm) and with low dispersal (no ballooning) that fell in pitfall traps in isolated plots. Thus, it contributed unique traits to roundabout communities. We finally used `kernel.originality` to estimate the functional originality of each species included in the dataset, finding low variability in originality values for species in both isolated and connected subplots. This is not unexpected since the pool of species found in urban environments are in general already filtered for a small diversity of traits (McKinney, 2006).

### Correlation with other indices

Using the dataset by Piano et al. (2020), we also explored the pairwise correlations among new indices in *BAT*, as well as the correlation with analogous indices calculated with *FD* (Laliberté et al., 2014) and *TPD* (Carmona et al., 2019). As far as we are aware, these are the only other R packages including functions for calculating richness, divergence, and regularity components of FD in a multidimensional space.

All community-level indices calculated with BAT were rather independent to one another (all |*r*|< 0.5; Figure 4a), and thus able to capture distinct facets of FD (Mouchet et al., 2010). The correlation between the two individual-level indices was also very low (*r*= 0.15; Figure 4b). In fact, contribution and originality are able to capture the contribution of each observation to the richness and divergence components of FD, respectively, allowing to map different components of functional rarity (Violle et al., 2017) at distinct scales of organizations (Carmona, de Bello, Sasaki, Uchida, & Pärtel, 2017).

On the other hand, the new indices are more highly correlated with similar indices in *FD* and *TPD* packages in terms of richness (Figure 4c) and divergence components (Figure 4d), whereas low correlation was found for Regularity (Figure 4e). These preliminary results emphasize the need to compare divergence among frameworks and properties of the existing indices, for example using simulations to test for different combinations of traits, composition of communities, and input parameters.

## Conclusions

By developing functions for estimating the primary facets of FD with *hypervolume*, we open up the potential to fully exploit the strengths of the Hutchinsonian niche concept (Hutchinson, 1957) in functional ecology research (Blonder, 2019). As in the case of other frameworks for estimating facets of FD, these new functions can be used to answer a large assortment of questions. Alpha and beta FD allow us to understand the richness of a functional hyperspace, and in which components multiple functional hyperspaces differ, namely the pairwise richness or replacement differences in term of functional traits. Contribution and originality allow to map which are the rare functional elements in the community. Therefore, these indices yield potential to help in targeting keystone species or those of high conservation concern (Carmona et al., 2017; Violle et al., 2017). Dispersion and evenness provide a quantification of the distribution of elements in the trait space enabling, for example, to compare whether multiple assemblages are differentially harmonious/unbalanced in their functional composition.

### CITATION

Researchers using these functions should cite this article and in addition can also cite the BAT package directly. Updated citation information can be obtained by typing in R the command `citation(‘BAT’)`. Parameters used for constructing hypervolumes have also to be specified, referring to the publications by Blonder et al. (2014, 2018).

## Acknowledgement

Special thanks to Benjamin Blonder for creating the hypervolume framework upon which our methods heavily rely, and to Elena Piano for providing data to test the new functions. Benjamin Blonder, Thomas Guillerme, and an anonymous referee provided useful suggestions throughout the review process.

## Data availability

Data and R scripts to generate the analyses are available in figshare (doi: 10.6084/m9.figshare.11708277). The package BAT is available through CRAN (https://cran.r-project.org/web/packages/BAT/), but it is also deposited in GitHub (https://github.com/cardosopmb/BAT).

